# Straightening FSC curves

**DOI:** 10.1101/2025.06.22.660909

**Authors:** Alexandre Urzhumtsev

## Abstract

The Fourier Shell Correlation (FSC) is traditionally calculated in resolution shells and plotted as a function of the inverse resolution. To ensure consistent statistics, these resolution shells should ideally contain an equal number of Fourier coefficients, at least approximately. This condition is naturally met when the shell boundaries are chosen uniformly in a scale based on the inverse cubic resolution. It has been observed that, in this scale, the FSC curves tend to exhibit local linearity. Using the new scale also zooms at the high-resolution shells allowing to extract values of interest more reliably.

**Synopsis:** Calculating and plotting FSC curves on a scale uniform in inverse cubic resolution (Å^−3^) has several useful features.

## 1. Introduction

Fourier Shell Correlation (FSC), computed as

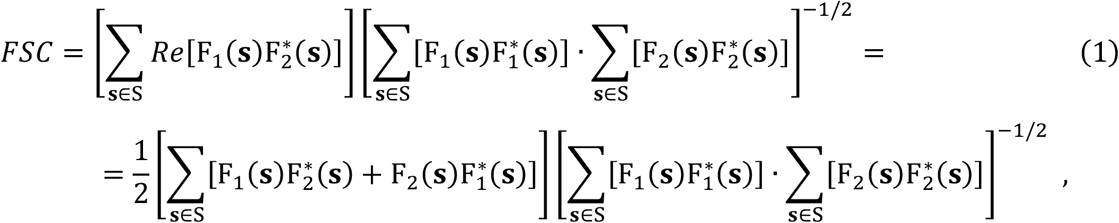

assesses the similarity between two sets of complex-valued Fourier coefficients, {F_1_(**s**)} and {F_2_(**s**)}, for a given set of indices, **s** ∈ S. In cryo electron microscopy (cryo-EM), the FSC analysis has been used since the 80ths (van Heel *et al*., 1982, Saxton & Baumeister, 1982; Harauz & van Heel, 1986; van Heel, 1987). In this context, the two sets of the Fourier coefficients may correspond to two maps or to a map and a model. Traditionally, FSC is used to compare two half-maps, which are independent 3D-reconstructions obtained from two complementary and randomly selected subsets, of nearly equal size, of the experimental data. For this analysis, FSC is traditionally calculated as a function of the inverse resolution, Å^−1^. The resulting curve generally has a sigmoid shape, with the FSC value equal to one for the inverse resolution close to zero *(i*.*e*., for lowest resolution) and decreasing, normally to zero, for high-resolution shells. In particular, one notes the lowest resolution *d*_*FSC*_ for which FSC drops below a chosen threshold value. This threshold indicates the limit beyond which the coefficients are considered as unreliable. Its values fixed at 0.5 (Böttcher *et al*., 1997; Beckman *et al*., 1997), 0.143 (Rosenthal & Henderson, 2003), or adjustable (“half-bit”, van Heel & Schatz, 2005) have been proposed and debated, with further discussions in the literature (*e*.*g*., Penczek, 2010; van Heel & Schatz, 2017) and online. Naturally, information contained in the FSC curves is not limited to a single *d*_*FSC*_ value, whatever its value may be, and “single-valued resolution criteria are often less relevant than the actual shape of an FSC curve” (Afanasyev *et al*., 2017).

Actually, this shape depends on the resolution scale which may be chosen differently depending on the specific problem. While this does not alter the information content of data, selecting an optimal scale may clarify its interpretation; in particular, adapting different scales to different tasks is implemented in the *Phenix* suite (Liebschner *et al*., 2019). For example, a uniform scale in Å^-2^ is optimal for calculating the so-called Wilson plot (Wilson, 1942) to estimate the value of the isotropic atomic displacement factor. A uniform scale in log(Å), where the number of reflections increases proportionally from one resolution shell to the next, has been shown useful both to analyze crystallographic *R*-factors (Urzhumtsev *et al*., 2009), and for cryo-EM studies (Stagg *et al*., 2014). Choosing the resolution shell boundaries uniformly in Å^−3^ ensures that these shells contain approximately equal number of coefficients. This latter feature appears particularly relevant for the calculation and interpretation of FSC curves.

In what follows, whenever possible, we use the more general notation, *d*^−3^, regardless of the particular units in which the resolution *d* is expressed.

## 2. Method

### 2.1. Features of the inverse cubic resolution scale

Since FSC curves are calculated over resolution shells, it is statistically appropriate for these shells to contain the same number of Fourier coefficients, at least approximately, provided that this number is statistically significant. This can be achieved by defining their boundaries uniformly with respect to the inverse cube of the resolution, *d*^−3^. Conversely, choosing resolution shells containing the same number of Fourier coefficients in three-dimensional space is equivalent to defining shell boundaries that are uniformly distributed in *d*^−3^.

In addition to the better sampling across a given spatial frequency range, another property is that using the scale in *d*^−3^ provides greater details in higher-resolution shells compared to the conventional scale in *d*^−1.^ This allows to locate more precisely the point of interest, *d*_*FSC*_, and this without increasing the total number of the resolution bins.

When computing the FSC as a function of *d*^−3^, it is natural to plot the resulted curves using the same scale. In the conventional *d*^−1^ scale, the FSC curves computed between two half-maps typically have a sigmoid shape. Therefore, certain common features can be also expected in the curves plotted with the modified scale.

### 2.2. Data used

To calculate, plot and analyze the FSC curves in the proposed scale *d*^−3^, we used deposited data from the EMDB (https://www.ebi.ac.uk/emdb/). The entries have been selected approximately at random, trying to illustrate a variety of resolutions. For each entry, the resolution *d*_*FSC*_ is reported by depositors as the point where the FSC curve reaches the values 0.143 or, alternatively, 0.5. In what follows, they are referred to as *d*_0.143_ and *d*_0.5_, respectively. Additional *d*_0.143_ and *d*_0.5_ values recalculated by the EMDB from the deposited data are also available. Although identifying the precise critical resolution was not the goal of this project, we used all these values as references, without delving into how they were computed or their exact interpretation.

### 2.3. Results presentation

The FSC curves were calculated in the resolution shells whose boundaries were defined uniformly in two different resolution scales, *d*^−1^ and *d*^−3^. In the current work, each curve extends to a resolution beyond which it has no structural significance (either oscillating around zero, or rising again after approaching zero, indicating residual noise coefficients). These upper resolution limits, *d*_high_, were chosen individually for each entry, and were set higher than the respective *d*_0.143_ values mentioned above (Table 1). For ease of comparison, all FSC curves shown here were computed using the same number of bins (resolution shells), arbitrary set to thirty, although other bin numbers, both smaller and larger, were tested in the course of this work. Varying the number of bins did not affect the observations discussed below. Figures 1 and 2 show the number of Fourier coefficients per bin, *N*_*coef*_, plotted as log_10_(*N*_*coef*_).

**Table 1.**
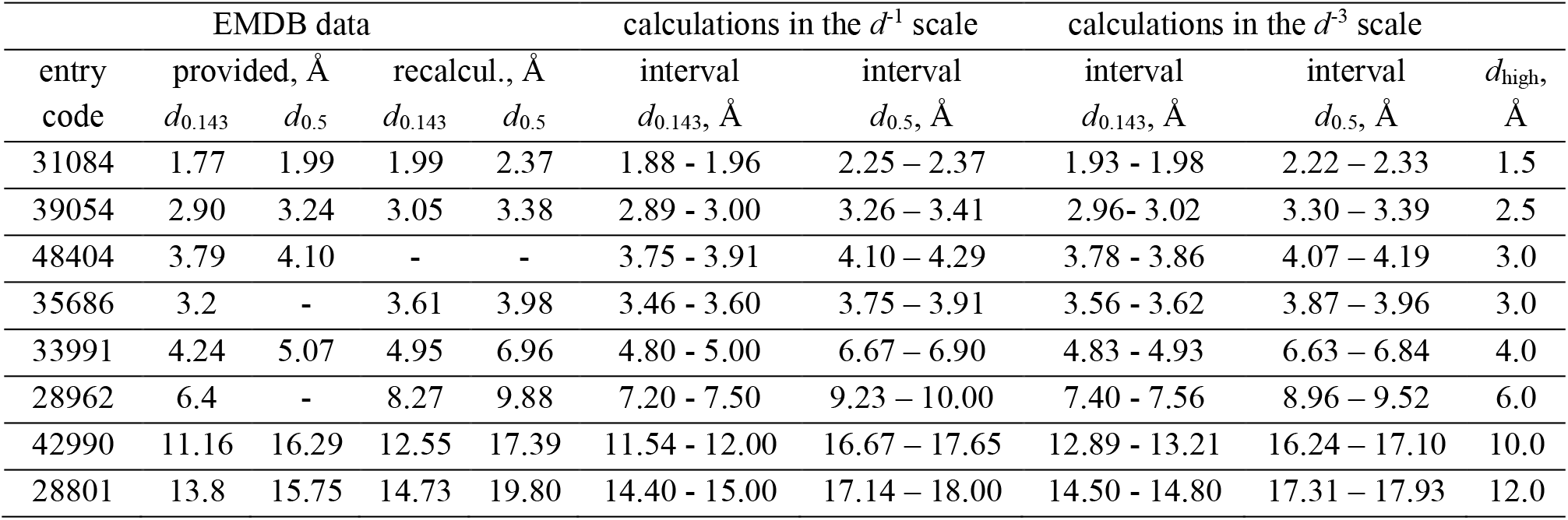
EMDB entries used. Columns 2^−3^ and 4-5 contain the *d*_0.143_ and *d*_0.5_ values provided by the depositors and independently recalculated by EMDB. Last column defines the resolution interval, 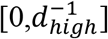 or 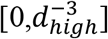, split into thirty bins with the boundaries chosen uniformly in the respective resolution scales and used for calculations. Columns 6 - 9 show the boundaries of the lowest-resolution bins where the FSC curves drop below the values 0.143 and 0.5, respectively. Number of the Fourier coefficients per bin is provided at Figures 1 and 2.

| entry code | EMDB data | | | | calculations in the $d^{-1}$ scale | | calculations in the $d^{-3}$ scale | | $d_{high}$, $\text{\AA}$ |
| --- | --- | --- | --- | --- | --- | --- | --- | --- | --- |
| | provided, $\text{\AA}$ | | recalcul., $\text{\AA}$ | | interval | interval | interval | interval | |
| | $d_{0.143}$ | $d_{0.5}$ | $d_{0.143}$ | $d_{0.5}$ | $d_{0.143}$, $\text{\AA}$ | $d_{0.5}$, $\text{\AA}$ | $d_{0.143}$, $\text{\AA}$ | $d_{0.5}$, $\text{\AA}$ | |
| 31084 | 1.77 | 1.99 | 1.99 | 2.37 | 1.88 - 1.96 | 2.25 - 2.37 | 1.93 - 1.98 | 2.22 - 2.33 | 1.5 |
| 39054 | 2.90 | 3.24 | 3.05 | 3.38 | 2.89 - 3.00 | 3.26 - 3.41 | 2.96 - 3.02 | 3.30 - 3.39 | 2.5 |
| 48404 | 3.79 | 4.10 | - | - | 3.75 - 3.91 | 4.10 - 4.29 | 3.78 - 3.86 | 4.07 - 4.19 | 3.0 |
| 35686 | 3.2 | - | 3.61 | 3.98 | 3.46 - 3.60 | 3.75 - 3.91 | 3.56 - 3.62 | 3.87 - 3.96 | 3.0 |
| 33991 | 4.24 | 5.07 | 4.95 | 6.96 | 4.80 - 5.00 | 6.67 - 6.90 | 4.83 - 4.93 | 6.63 - 6.84 | 4.0 |
| 28962 | 6.4 | - | 8.27 | 9.88 | 7.20 - 7.50 | 9.23 - 10.00 | 7.40 - 7.56 | 8.96 - 9.52 | 6.0 |
| 42990 | 11.16 | 16.29 | 12.55 | 17.39 | 11.54 - 12.00 | 16.67 - 17.65 | 12.89 - 13.21 | 16.24 - 17.10 | 10.0 |
| 28801 | 13.8 | 15.75 | 14.73 | 19.80 | 14.40 - 15.00 | 17.14 - 18.00 | 14.50 - 14.80 | 17.31 - 17.93 | 12.0 |

**Figure 1.**
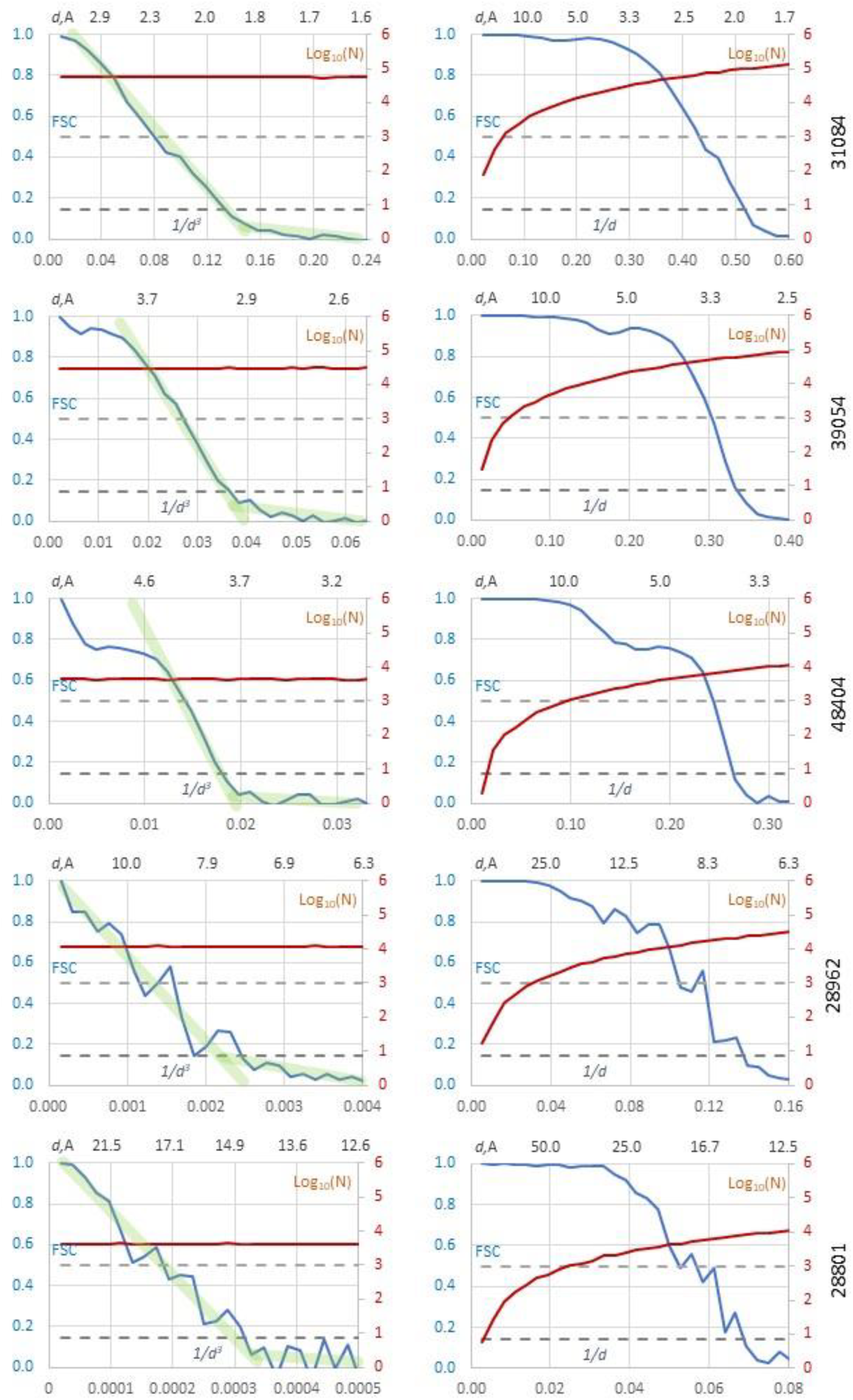
FSC curves. FSC curves (in blue) were calculated in resolution shells with boundaries chosen uniformly in the Å^-3^ (left column) and Å^−1^ (right column) scales. The upper axis shows the resolution recalculated in Å. The marron curves represent the log_10_ of the corresponding number of Fourier coefficients; note their pronounced variation for the Å^−1^ scale. Dashed lines indicate the FSC values of 0.143 and 0.5. The EMDB entry code for each dataset is shown nearby.

**Figure 2.**
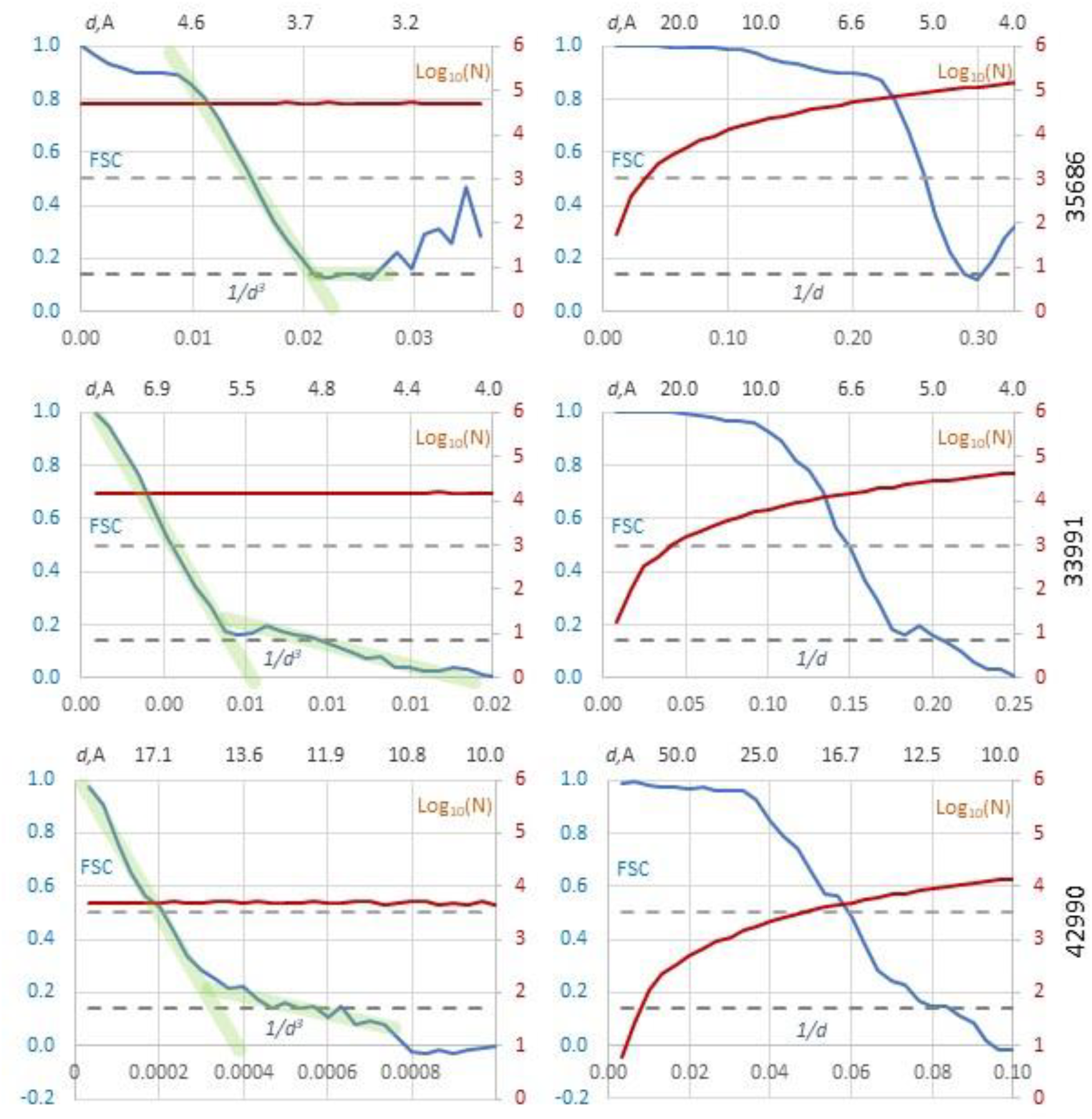
FSC curves. FSC curves (in blue) were calculated in resolution shells with boundaries chosen uniformly in the Å^-3^ (left column) and Å^−1^ (right column) scales. The upper axis shows the resolution recalculated in Å. The marron curves represent the log_10_ of the corresponding number of Fourier coefficients; note their pronounced variation for the Å^−1^ scale. Dashed lines indicate the FSC values of 0.143 and 0.5. The EMDB entry code for each dataset is shown nearby.

The diagrams show the calculated FSC values as a function of the resolution of the respective shells. For each calculation, we noted the interval where the FSC curve crosses the prescribed threshold values of 0.143 and 0.5 (Table 1). No interpolation within these intervals was performed to further refine the corresponding *d*_0.143_ and *d*_0.5_ values.

Another reference resolution, denoted here as *d*_*ss*_, corresponds to the interval around 5 – 6 Å where the FSC curves typically exhibit a small local minimum and maximum. This feature is generally attributed to the presence of the macromolecular secondary structure elements such as *α*-helices and *β*-sheets.

## 3. Results

Table 1 lists the EMDB entries analyzed in this article and illustrated by Figures, along with key characteristics of the corresponding data sets. As expected, for the same number of intervals, the calculations performed using the *d*^−3^ scale yield intervals containing *d*_0.143_ roughly half the width compared to those using the *d*^−1^ scale (columns 6 and 8), while the width of the intervals for *d*_0.5_ remain comparable (columns 7 and 9). Overall, these intervals are in a reasonable agreement with the *d*_0.143_ and *d*_0.5_ estimates provided by EMDB.

Figure 1 illustrates FSC curves for data sets spanning a range of resolutions, from high to very low. A pronounced variation in the number of coefficients per bin is observed when using the *d*^−1^ scale whereas this number remains nearly constant for the calculations in the *d*^−3^ scale. This consistency is the principal argument for adopting the new scale.

The non-linear rescaling from *d*^−1^ to *d*^−3^ induces a corresponding non-linear transformation of the FSC curves, changing their shape from sigmoid-like to rather L-shaped. The slopes of the linear segments in this approximated shape highlights a difference in characteristics of the Fourier coefficients in different resolution regions, and, together with the coordinates of their intersection point, may potentially serve as an indicator of map quality, similar to the FSC ‘knee’ for the curves in the *d*^−1^ scale pointed out by Afanasyev *et al*. (2017).

For high- and medium-resolution data (EMDB entries 31084, 39054, 48404), the FSC curves, starting from *d*_*ss*_ (5 - 6 Å), follow an approximately straight line up to a certain point, beyond which they oscillate around another nearly straight line with a much smaller slope. For lower resolution data (EMDB entries 28962 and 28801) the FSC curve plotted in the *d*^−3^ retains its L-shape, were the ‘information-containing’ part of the curve again oscillates around a straight line. However, these oscillations are larger than for the previous entries, even when the number of coefficients per bin remains relatively high (compare these numbers for entries 28962 and 48404).

Figure 2 illustrates additional examples. For a high-resolution case (EMDB entry 35686), the right-hand of the FSC curve oscillating around a nearly horizontal straight-line segment indicating unconfident Fourier coefficients whether their values are slightly higher or lower 0.143. In a medium resolution example (EMDB entry 33991), the two parts of the FSC curve again oscillate around two straight lines, in this case being separated by the local minimum at *d*_*ss*_. For low-resolution data (EMDB entry 42990), the right part of the curve also oscillated around the 0.143 level, similar to entry 35686. A smoother transition between the two straight-line segments may be explained, at least partially, by the relatively small number of the Fourier coefficients per resolution shell for this low-resolution case.

Working in the Å^−3^ scale allows for a more detailed examination of the FSC curve at higher resolutions while keeping the same total number of bins. Conversely, this compresses the low-resolution Fourier coefficients into fewer bins, which may obscure FSC information in the region around the *d*_*ss*_ resolution, another area of possible interest. To address this problem, such features may be analyzed separately by selecting a standard resolution interval, for example that up to the 4.5 Å resolution, which corresponds to approximately 0 ≤ *d*^−3^ ≤ 0.011 Å^−3^ (Fig. 3).

**Figure 3.**
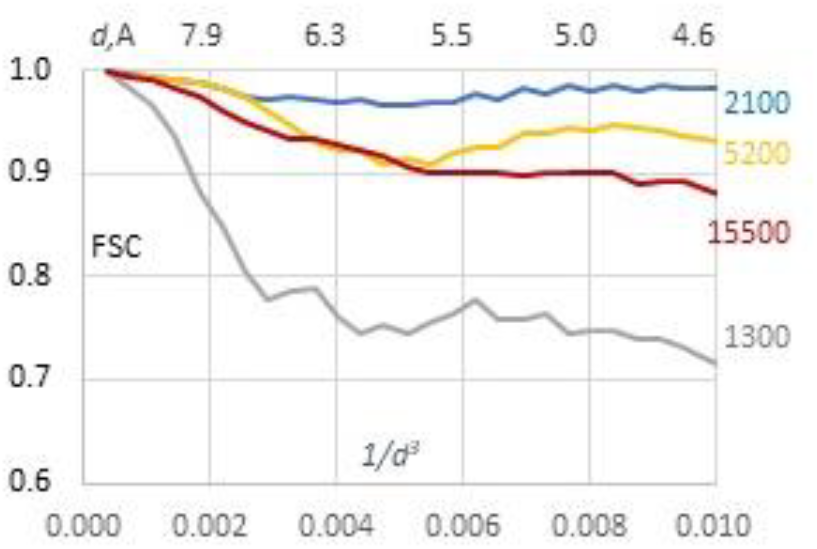
FSC curves. FSC curves calculated in the resolution shells with the boundaries chosen uniformly in the Å^−3^ scale up to the resolution 4.5 Å (0.011 Å^−3^). Upper resolution scale is reclculated in Å. Mean number of the Fourier coefficients per resolution shell is given nearby the curves.

## 4. Discussion

Calculating FSC curves between two sets of Fourier coefficients in the Å^−3^ scale yields a balanced distribution of the coefficients per shell and thus more reliable statistical values than using the conventional scale in Å^−1^. From a mathematical standpoint, it renders the statistical analysis more robust and consistent, which alone provides a strong rationale for using the new scale. This improvement can be particularly important when comparing two half-maps, as improved statistical handling may result in a more reliable interpretation of FSC curves.

In this new scale, FSC curves also exhibit a characteristic shape, as observed previously for the curves plotted in Å^−1^. Analyzing the overall shape of the FSC curve can provide more meaningful insights into data quality than focusing solely on a specific threshold value, definition of which with a high precision may be overly theoretical and sometimes leading to a controversy in practical FSC-based resolution estimations.

For the FSC curves plotted in the Å^−3^ scale, their oscillation around straight lines may offer a simpler way to interpret them than the traditional sigmoid shapes seen in the conventional Å^−1^ scale, although this raises some questions. For instance, could the parameters of the intersection point of these straight segments help in estimating the threshold beyond which Fourier coefficients start missing confident and useful information? Does the slope of these straight segments have any statistical significance? Analysis of these and related questions, including accurate and unbiased determination of *d*_0.143_ or other critical values, is beyond the scope of this article, which primarily aims to demonstrate the statistical benefits of using a uniform scale in Å^−3^.

In summary, the principal advantage of computing the FSC curve in a uniform scale in Å^−3^ is the improved statistical consistency, similar to calculation of the crystallographic *R*-factors as a function of resolution. Second, automatically, this zooms the curves at the higher resolution region without increasing the total number of resolution intervals compared to the conventional Å^−1^ scale. Finally, for half-map analysis, switching to this new scale transforms the typical sigmoid-type curves into a simpler form, where FSC curves tend to follow, or oscillate around, two straight segments. Such clearer shapes may provide more insight into map quality and help mitigate common ambiguities in determining the critical resolution.

Obviously, this modification of the resolution scale entails the non-negligible inconvenience of departing from established habits and working with a less familiar curve shape. However, this can be regarded much like adopting a new version of a conventional software: initially unfamiliar, but likely to become routine practice once its advantages are recognized.

## Acknowledment

The author thanks P.V. Afonine and B.P. Klaholz for help with the data and extremely stimulating discussions, and acknowledges Instruct-ERIC and the French Infrastructure for Integrated Structural Biology FRISBI [ANR^−1^0-INBS-05].

